# Patient-derived Cellular Models of Polygenic Scores: Application to Long QT Syndrome

**DOI:** 10.1101/2025.08.05.667271

**Authors:** Devyn M. Mitchell, Matthew J. O’Neill, Loren Vanags, Nidhi Patel, Theresa Strickland, Giovanni Davagustto, M. Benjamin Shoemaker, Joe Elie Salem, Brett M. Kroncke

## Abstract

Polygenic scores (PGS) have emerged as important modifiers of disease risk, drug response, and rare variant penetrance, but the biological mechanisms underlying these associations remain poorly understood. To experimentally investigate these relationships, we derived induced pluripotent stem cell-derived cardiomyocytes (iPSC-CMs) from individuals at the 1st, 50th, and 99th percentiles of a genome-wide QT interval polygenic score (QT-PGS). We examined baseline repolarization, drug response to the hERG inhibitor E4031, and penetrance of two KCNH2 variants (p.Arg148Trp and p.Arg823Trp) in these cellular models. While baseline field potential durations (FPDs) did not differ across PGS levels, high-PGS iPSC-CMs showed exaggerated prolongation in response to E4031 and increased phenotypic expression of both KCNH2 variants. These findings provide the first experimental demonstration that polygenic background can shape cardiac electrophysiologic phenotypes and modulate the functional impact of both pharmacologic and genetic perturbations. This work establishes a scalable platform for mechanistic studies of polygenic risk.

## Introduction

Historically, cardiovascular genetics have focused on rare variants with large effect size, such as loss-of-function variants in ion channels associated with congenital Long QT Syndrome (cLQTS). More recently, BioBank scale studies have revealed a cumulative influence of common variant burden on traditional ‘Mendelian’ disease. These polygenic scores (PGS) have been associated with variability across many traits, and have been shown to modulate drug response and rare variant penetrance in observational studies (*1, 2*). To date, the underlying mechanisms remain elusive, especially hindered by the absence of controlled experimental models. The possibility of studying patient-derived cell-lines harboring the extremes of estimated QT polygenic variability represents a novel approach to experimentally probe this burden for the first time.

## Results/Discussion

Accordingly, we derived induced pluripotent stem cell cardiomyocytes (iPSC-CMs) from individuals at the extremes of a QT-PGS derived from 1,107,630 single nucleotide polymorphisms (1^st^, 50^th^, and 99^th^ percentile; **Figure A**) (*2, 3*). Three individuals were identified and consented, after which iPSC-CMs were derived from their peripheral blood mononuclear cells using previously described techniques, under approval of IRB 211014 (*4*). We exploited this unique resource to address three lines of inquiry: 1) can a QT-PGS extreme approximate a monogenic cLQTS phenotype? 2) do QT-PGS extremes modulate drug-induced prolongation of ventricular repolarization? 3) do QT-PGS extremes affect traditional ‘Mendelian’ rare variant penetrance? Specifically, we interrogated rare variants in *KCNH2*, the gene encoding the rapidly inactivating cardiac potassium channel (hERG) associated with ∼40% of cLQTS (OMIM: 613688), and a frequent rate-limiting target of drug candidates.

**Figure 1.**
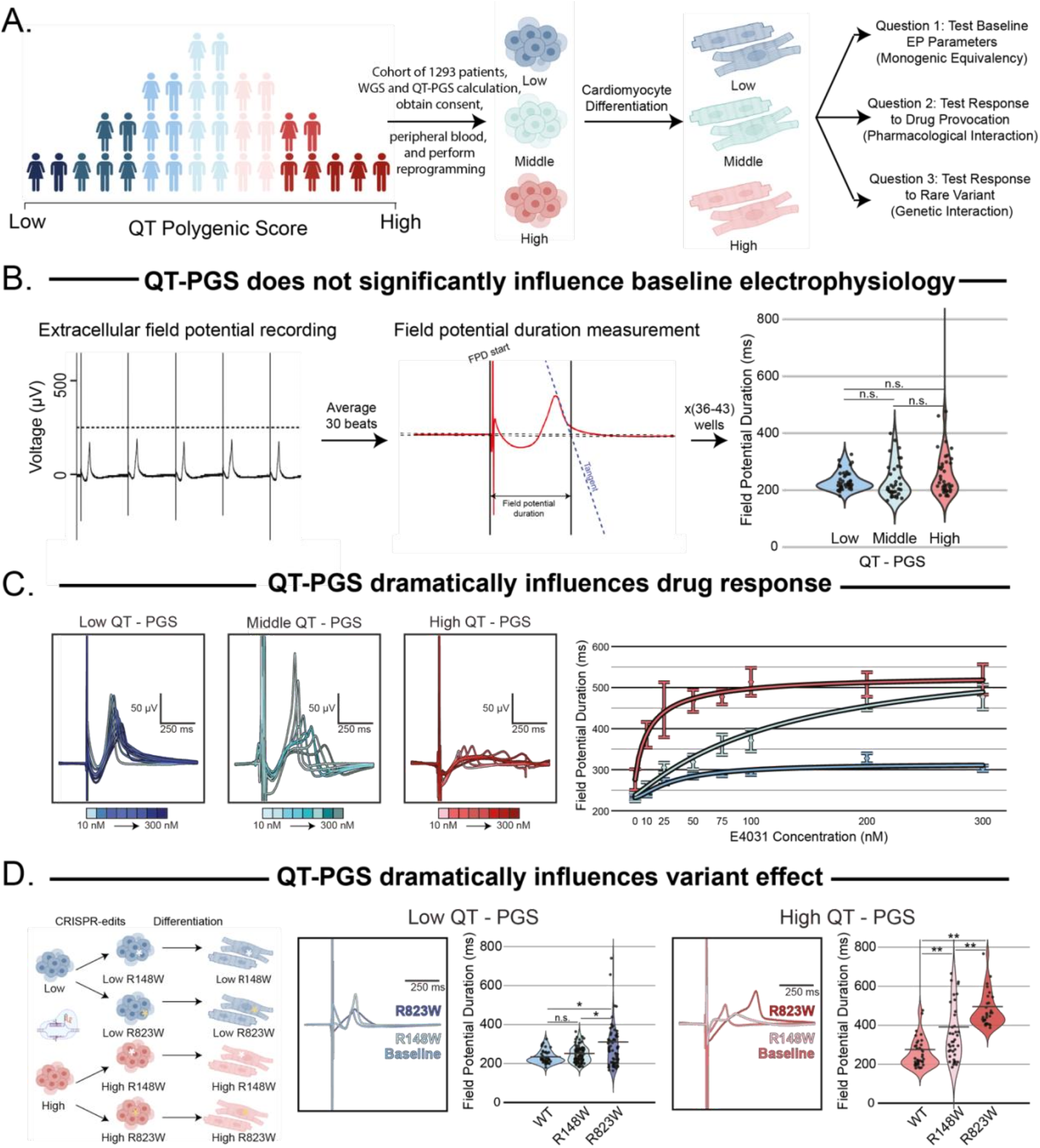
**A)** Schematic overview of project. Patients at extremes of the QTc interval Polygenic Scores were identified (QT-PRS). Induced pluripotent stem cell-cardiomyocytes (iPSC-CMs) were derived from peripheral blood mononuclear cells, from which three key questions were queried. **B)** iPSC-CMs were studied using the Nanion CardioExcyte system. Example well tracings over time are displayed. Field potential durations were derived from an average of 30 beats per well. Violin plots of FPD compare baseline values (Wilcoxon rank sum test, False Discovery Rate [FDR] < 0.05). **C)** Field Potential Duration (FPD) traces were recorded across the three lines under increasing titration of E4031, a chemical tool that inhibits hERG. Summary data of average and standard error FPD measurements across each cell line under increasing doses of E4031. **D)** Schematic of CRISPR-edited iPSC-CMs. Single well representative trace of EFP for wild type lines compared to those harboring edits of p.Arg148Trp and p.Arg823Trp. Right, comparison of mean and standard error of the mean for each condition (^*^ indicates statistically significant under FDR < 0.05; ^**^ < 0.005).

First, we studied cellular proxies of ventricular repolarization at baseline across low-, middle-, and high QT-PGS iPSC-CMs. We recorded extracellular field potentials (EFP) and then derived Field Potential Durations (FPDs), which are cellular measurements of action potential duration, mirroring QTc measured clinically (**Figure B**). These recordings were taken across 36-43 quality control passing wells per line (monolayers of ∼50,000 hiPSC-CMs/well), using the Nanion CardioExcyte platform (Munich, Germany; **Figure B**). All cells were paced at 1 Hz with minor variability, corrected by Fridericia’s formula as necessary. Previous studies have shown that certain extremes of PGS may reflect monogenic equivalency among Tier 1 conditions in BioBanks (*5*). In our experimental study, we observed no significant difference in FPDs among iPSC-CM monolayers derived from low, middle, and high QT-PGS individuals at baseline (Wilcoxon rank-sum test, False Discovery Rate < 0.05).

Second, we studied how PGS extremes would affect responsiveness to the QT-prolonging compound E4031. Former studies have shown that common variant burden may modulate QT-prolonging drug effects (*2*). After E4031 administration, we observed a non-linear increase in FPD, with increased susceptibility among the high vs middle vs low line across progressive drug titration (average 39 wells per condition, minimum 25; **Figure C**).

Third, we probed whether extremes of QT-PGS would affect penetrance of rare variants with incomplete penetrance for cLQTS. A previous study has shown that common variant burden may modulate rare variant penetrance in human BioBank studies (*1*). We studied two *KCNH2* variants with different degrees of loss-of-function: p.Arg148Trp (ClinVar: 67505, Conflicting Interpretations; mean QTc 441 ms, n = 27, variantbrowser.org) and p.Arg823Trp (ClinVar: 67402, Pathogenic/Likely Pathogenic; mean QTc 487 ms, n = 15, variantbrowser.org) **(Figure D)**. Accordingly, we introduced each variant into the low- and high-PGS lines using CRISPR-Cas9 editing, as previously described by our lab (**Figure D**). We observed that the low-PGS line attenuated the less penetrant variant p.Arg148Trp, but that the more highly penetrant variant p.Arg823Trp significantly affected FPD (**Figure D**). In contrast, the high PGS line showed similar baseline FPD, but significantly increased FPDs when harboring either variant (**Figure D**). Collectively, this work presents a proof of concept to interrogate BioBank-scale observations of QT-PGS experimentally for the first time. We believe this bold approach to describing and eventually interrogating QT-PGS mechanisms may lead to enhanced risk stratification and discovery opportunities.

## Materials and Methods

CRISPR-Cas9 Gene Editing. To introduce KCNH2 variants (c.442C>T [p.Arg148Trp] and c.2467C>T [p.Arg823Trp]), CRISPR-Cas9 genome editing was performed on iPSC lines from low- and high-QT-PGS individuals. Guide RNAs were designed using the online webserver CRISPOR.org and cloned into the pSpCas9(BB)-2A-GFP (PX458) vector (Addgene), which co-expresses Cas9 and green fluorescent protein (GFP). A single-stranded DNA repair template was designed to introduce the desired variant. In both cases, the protospacer-adjacent motif (PAM) site overlapped with the target mutation and could not be modified independently; therefore, a synonymous nucleotide polymorphism was incorporated into the repair template to prevent re-cleavage by Cas9. Repair template for facilitating homology directed repair included a SNP in the repair template using a silent mutation to preserve the amino acid sequence (lowercase and bolded): pArg148Trp: 5’-GTG ATG GAG AAG GAC ATG GTG GGG TCC CCG GC**a** CAT GAC ACC AAC CAC **t**GG GGC CCC CCC ACC AGC TGG CTG GCC CCA GGT AAG TGT ACT TGC TTT AGG G -3’, p.Arg823Trp: 5’-GC TCC CTG TCC TCT CCA TGG CCT CCA GGG AAG AAT GAC ATC TTT GGG GAG CCT CTG AAC CTG TAT GCA AGG CCT GGC AAG TCG AA**t** GGG GAT GTG tGG GCC CTC ACC TAC TGT GAC CTA CAC AAG-3’. iPSCs were grown to 70–80% confluency and electroporated with the modified PX458vector and repair template using the Neon Transfection System (settings: 1650 V, 10 ms pulse width, 3 pulses). Following electroporation, GFP positive cells were isolated by flow cytometry, cultured for a least 5 days, and subsequently screened and confirmed by bidirectional Sanger sequencing. Edited clones were expanded, cryopreserved, and revalidated for pluripotency and karyotypic stability before differentiation into iPSC-CMs.

Cardiac Differentiation of iPSCs. iPSCs were expanded in mTeSR1 medium for 4–5 days until reaching approximately 80% confluency. Differentiation into cardiomyocytes (iPSC-CMs) was initiated by switching to cardiac differentiation medium consisting of RPMI 1640 basal medium (catalog no. 11875; Life Technologies, Carlsbad, CA), supplemented with 2% B-27 minus insulin (catalog no. A1895601; Life Technologies), and 1% penicillin–streptomycin (Life Technologies). From days 0 to 2 of differentiation, the medium was supplemented with 6 μM CHIR99021 (LC Laboratories, Woburn, MA). From days 3 to 5, it was supplemented with 5 μM IWR-1 (Sigma-Aldrich, St. Louis, MO). Media was exchanged every other day. Metabolic selection was performed from days 13 to 19 using glucose-free RPMI 1640 (catalog no. 11879; Life Technologies), supplemented with 2% B-27 minus insulin and 1% penicillin–streptomycin. From day 19 onward, cells were maintained in cardiomyocyte maintenance medium (RPMI 1640 with 2% B-27 supplement including insulin; Life Technologies), with media changes every other day. Differentiated iPSC-CMs were validated by immunofluorescence staining for cardiac markers, including alpha-actinin (Sigma-Aldrich) and myosin light chain-2V (ProteinTech Group, Rosemont, IL) (Figure S2).

Optical Pacing Vector Transduction and Cell Replating. To ensure pace rate at or near 1Hz, iPSC-CMs were transduced between days 28 and 30 post-differentiation with an adeno-associated virus (AAV) vector encoding channelrhodopsin-2: AAV1.CAG.hChR2(H134R)-mCherry.WPRE.SV40 (Addgene plasmid no. 20938M) at a multiplicity of infection of 6500 genome copies per cell. At day 30, iPSC-CMs were dissociated using TrypLE Express (Life Technologies) and replated at a density of 50,000 cells per well in 96-well CardioExcyte sensor plates (Nanion Technologies, Munich, Germany) precoated with 50 μL of Matrigel diluted 1:100 in RPMI 1640. Cells were allowed to recover and form monolayers, and electrophysiological studies were conducted between days 38 and 42 post-differentiation. In cases where cells exhibited spontaneous beating at approximately 1 Hz, optical pacing was not applied to avoid interference.

Electrophysiological Recordings. Extracellular field potentials (EFPs) and impedance signals were recorded using the CardioExcyte 96 platform (Nanion Technologies) 8–12 days after replating (corresponding to days 38–42 post-differentiation). Plates were maintained at 37°C, 80% humidity, and 5% CO2 in a controlled incubator integrated with the system. For optical pacing, a custom stimulating optical lid (Nanion Technologies) was used to deliver 470 nm wavelength light pulses across all wells. Cells were paced at 1 Hz with 5 ms pulse durations at 30 mA power. Recordings consisted of 20–30 second sweeps acquired at a sampling rate of 10 kHz (0.1 ms interval). Optical pacing was selected over electrical stimulation to minimize artifacts in EFP signals, allowing for cleaner repolarization measurements. Field potential duration (FPD), a cellular proxy for action potential duration and clinical QT interval, was calculated from the EFP waveforms. FPD values were corrected for beating rate variability using Fridericia’s formula: FPD = original FPD /(RR interval)^1/3^, where RR is the interval between consecutive beats, called FPD throughout. Baseline recordings included 36-43 quality-controlled cells per iPSC line, filtered for at least 2Ω impedance and 10uV signal stability reflecting monolayer integrity.

Drug Treatment and Response Assessment. To evaluate drug-induced repolarization changes, iPSC-CM monolayers were exposed to the hERG inhibitor E4031 (1 mM stock concentration). Drug was administered in a progressive titration protocol, with concentrations incrementally increased, waiting 1 hour while monitoring EFP responses then recording. Recordings were performed under the same conditions as baseline measurements, with an average of 39 cells per condition (minimum 25 cells) across low-, middle-, and high-QT-PGS lines.

## References

1. A. C. Fahed et al., Polygenic background modifies penetrance of monogenic variants for tier 1 genomic conditions. Nat Commun 11, 3635 (2020).

2. M. C. Lancaster et al., A Polygenic Predictor of Baseline QTc is Associated With SotalolInduced QT Prolongation. Circulation 150, 1984–1986 (2024).

3. V. Nauffal et al., Monogenic and Polygenic Contributions to QTc Prolongation in the Population. Circulation 145, 1524–1533 (2022).

4. D. Mitchell et al., Generation of human induced pluripotent stem cell (hiPSC) lines from patients with extreme high and low polygenic scores for QT interval. Stem Cell Res 84, 103691 (2025).

5. A. V. Khera et al., Genome-wide polygenic scores for common diseases identify individuals with risk equivalent to monogenic mutations. Nat Genet 50, 1219–1224 (2018).

